# Grid pattern development, distortions and topological defects may depend on distributed anchoring

**DOI:** 10.1101/830158

**Authors:** Maria Mørreaunet, Martin Hägglund

## Abstract

The firing pattern of grid cells in rats has been shown to exhibit elastic distortions that compresses and shears the pattern and suggests that the grid is locally anchored. Anchoring points may need to be learned to account for different environments. We recorded grid cells in animals encountering a novel environment. The grid pattern was not stable but moved between the first few sessions predicted by the animals running behavior. Using a learning continuous attractor network model, we show that learning distributed anchoring points may lead to such grid field movement as well as previously observed shearing and compression distortions. The model further predicted topological defects comprising a pentagonal/heptagonal break in the pattern. Grids recorded in large environments were shown to exhibit such topological defects. Taken together, the final pattern may be a compromise between local network attractor states driven by self-motion signals and distributed anchoring inputs from place cells.

## Introduction

Cortex may represent variables in discrete or continuous low dimensional attractor states. Although point attractors may be stably learned, for instance by Hopfield networks(Hopfield 1982), continuous attractors networks (CANs) are notably difficult to balance as small perturbations will lead them to wander away from the proper state. Thus, CANs need to monitor and reset their state either by lower level sensory inputs or by learned associations with representations found in higher cortical levels.

Grid cells in the medial entorhinal cortex fire in a periodic hexagonal pattern which display translational symmetry across cells suggesting they may use low dimensional continuous attractor dynamics to function as a path integrator by taking self-movement signals as inputs to update the animal’s representation of position in allocentric space(Fuhs and Touretzky 2006; Hafting et al. 2005; McNaughton et al. 2006; Yoon et al. 2013). The grid pattern is not perfectly hexagonal but displays elastic distortions that shears the grid along the orientation of a square environment(Stensola et al. 2015). Moreover, local elastic distortions have been described that stems from an interaction with the borders of the environment, compressing the grid along the axes of the walls(Hagglund et al. 2019). Crystalline materials may undergo similar elastic distortions to a certain degree, after which increased shear strain is released by slippage of a grain boundary, or nucleation of topological defects such as dislocations. Whether strong local elastic distortions may lead to similar topological defects in the grid pattern is unknown, but they would provide strong evidence of anchoring mechanics.

A concern for any system that relies solely on internal locomotor cues to update a position estimate is sensitivity to noise. Incomplete or imperfect sensory input may impart drift to the representation and undermine the possibility to derive stable spatial information from it over time. This caveat could be mitigated by a resetting mechanism that would ensure the systems long-term stability. Thus, to be useful as a spatial reference the grid needs to anchor to its environment. Anchoring has been suggested as a process where stable features of the environment provides a reference frame that will reset the position estimate at these various locations(Campbell et al. 2018; Dordek et al. 2016; Mulas, Waniek, and Conradt 2016; Ocko et al. 2018). However, compression distortions lead to a gradual contortion of the pattern across the environment which may necessitate resetting to occur at multiple locations distal to local cues and borders to keep the pattern stable. Anchoring further implies learning which has been observed where the grid has been locally altered by local changes in the environment(Boccara et al. 2019; Sanguinetti-Scheck 2019; Wernle et al. 2018). Thus, effects of anchoring in an open field may be most conspicuous during initial encounters with a novel environment where the pattern may depend more strongly on plastic interactions with the environmental layout. The grid has been shown to get increasingly hexagonal over the first few sessions after being subjected to a novel environment(Barry et al. 2012), possibly as a process of increasingly acquired anchoring points.

Here we demonstrate how the grid pattern develops in relation to behavior when subjected to a novel environment, how anchoring may explain details of the development as well as previously described distortions of the grid, and how anchoring may affect the topology of the grid pattern.

## Results

### GRID FIELDS MOVE OPPOSITE TO THE RUNNING DIRECTION DURING NOVELTY

To investigate the impact of learning on anchoring and the form of the grid we implanted rats with multi-tetrode devices and recorded grid cells in the medial entorhinal cortex. Animals foraged for rewards in a square 1.5 m box, a 1.8 m equilateral triangle or a 2.4 m equilateral triangle for a novel session followed by several subsequent sessions in the same box between 5 h to 3 days later (5 animals in 9 novelty exposures, 40 grid cells, 2-6 grid cells recorded simultaneously in each experiment). Cells that had a gridness score(Langston et al. 2010) above that of the 95^th^ percentile of a shuffled distribution was chosen, if this were true for at least one of the sessions. Thus, some cells in the analyses did not qualify as a grid cell in the initial session. Only cells with grid spacing smaller than 75 cm was chosen to improve the resolution of the local analyses below.

Local changes in the grid were studied by subdividing firing rate maps into 9 equal squares for the square environments (Figure 1A), 9 equilateral triangles for the large triangular environment and 4 equilateral triangles for the smaller triangle, leading to subdivisions of 0.25 m^2^, 0.28 m^2^ and 0.45 m^2^ respectively. For each cell, each subdivision from session 1 was cross-correlated with that from session 2 (Figure 1A, blue vs red respectively). Spatial differences in the grid between session 1 and 2 was measured from the offset of the peak closest to the middle of the cross-correlogram in each subdivision, if the peak was higher than 0.4 r and the peak had not moved more than 25% of the grid spacing of the cell. Grid fields were not stable between the 1^st^ and 2^nd^ session but had shifted a mean of 5.1 cm (sd = 3.2 cm, Figure 1B). There were no differences in the magnitude of the shift in the corners, along the walls nor in the middle subdivisions of the experiments in the square environment (mean magnitude 5.0 cm, sd = 2.9; 5.1 cm, sd = 3.6; and 5.4 cm, sd 2.7, in the mid, corner and wall compartments respectively, Wilcoxon’s rank sum test of the magnitude in the mid vs corner compartments, p = 0.96, mid vs wall, p = 0.35, and corner vs wall compartments, p = 0.21).

**Figure 1.**
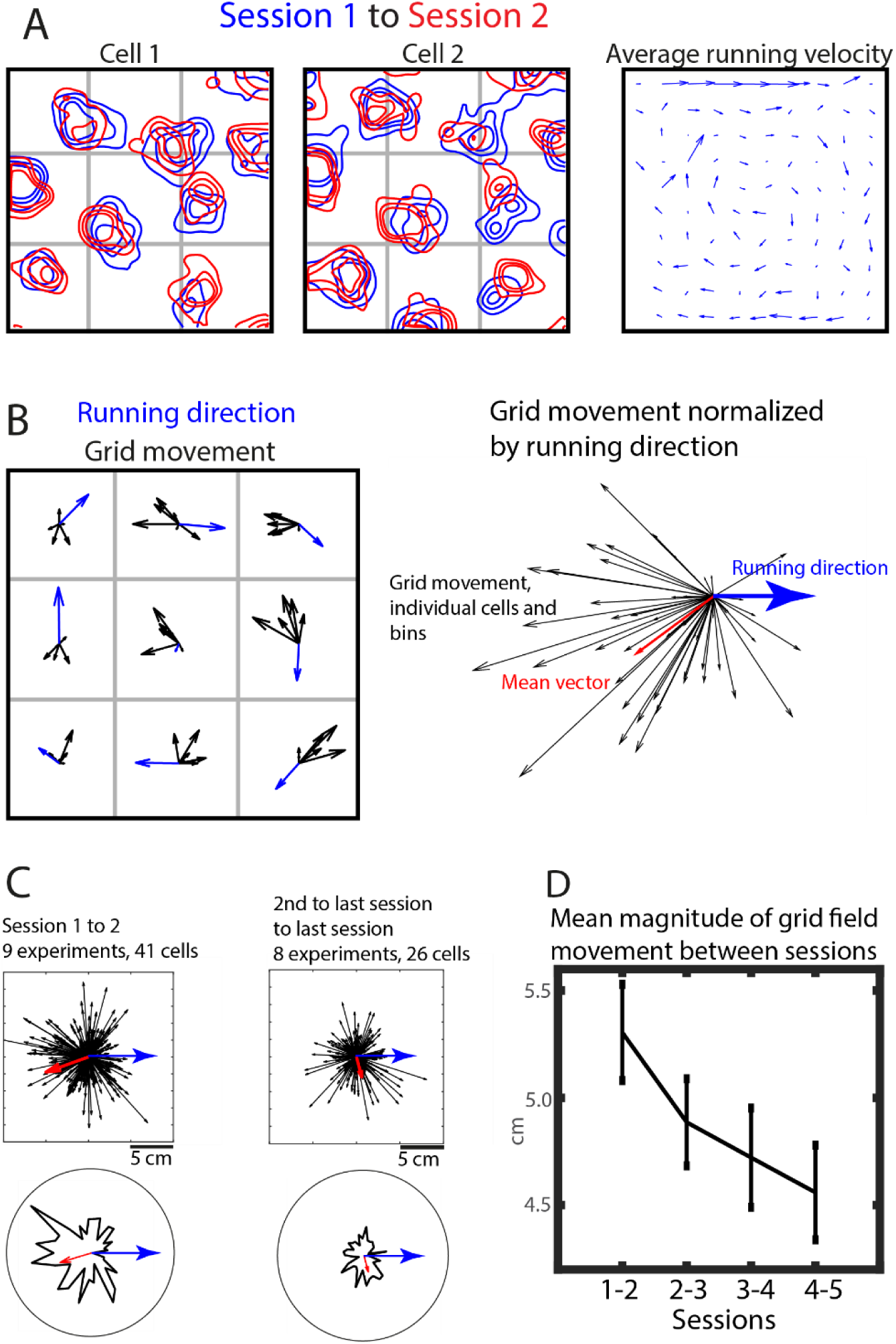
Individual grid fields move oppositely to the average running velocity during novelty. **(A)** Two example cells recording during novelty (blue) and a subsequent session (red) are shown together with the average running velocity (right). **(B)** The cross-correlation between sub compartments in session 1 and 2 shows how the grid moves locally (left, black arrows, 6 cells) displayed in relation to the average running velocity (blue arrows). Normalizing the grid movement to the running direction in this example shows that most shifts occur in the opposite direction of the average running direction (right). **(C)** Pooling the shifts normalized to average running direction shows that the shifts from 40 grid cells from 9 experiments run in the opposite direction of the behavior (left). Black arrows denote field shifts, the red arrow displays the mean vector and the blue running direction. The circular histogram is divided into 10° bins and displays counts weighted by shift magnitude (bottom). The 2^nd^ to last vs last session grid movement was smaller in magnitude and did not shift in any significant direction (C). **(D)** The mean magnitude of movement dropped during subsequent exposure to the environment (mean and SEM).

Across animals, the grid did not shift in any coherent pattern (N = 303 subdivision measurements, mean resultant vector length (MVL) = 0.052, weighted circular mean direction = 21°, Rayleigh’s test for circular non-uniformity, p = 0.18). However, when correcting for the behavior within each subdivision by rotating the grid shift direction according to the prevailing running direction in that subdivision from the first session, the shift of the grid was significantly distributed oppositely to the average local running direction (Figure 1B right shows example and Figure 1C, left, shows the entire data set; MVL = 0.17, weighted circular mean direction = 197°, Rayleigh’s test for circular non-uniformity, p = 0.031, weighted circular V-test for non-uniformity around the specific direction 180° from the average running direction (V-test_180_), v = 179, p = 9.4 × 10^−15^). Using a subset of the same cells recorded at later sessions demonstrated that the effect was due to novelty (Figure 2C, right; 26 grid cells comparing sessions 4 and 5, 7 and 8, 3 and 4, 5 and 6, 2 and 3, 5 and 6 and finally 5 and 6 for the respective experiments, MVL = 0.089, weighted circular mean direction after rotating according to running direction = 284°, V-test_180_, v = −13, p = 0.78). Although the direction of the shift in relation to the running direction was significant, the resultant mean vector length was short, suggesting that the grid may move locally also in other directions, possibly to consolidate a less regular initial pattern (Barry et al. 2012). The movement of the grid fields opposite to behavior was significant between session 1 and 2 and between session 2 and 3 (MVL = 0.10, V-test_180_, V = 58, p = 0.0036) after which any movement was statistically undetectable by these means (Figure 1D, session 3 to 4, MVL 0.07, V-test_180_, V = −8.3, p = 0.67).

**Figure 2.**
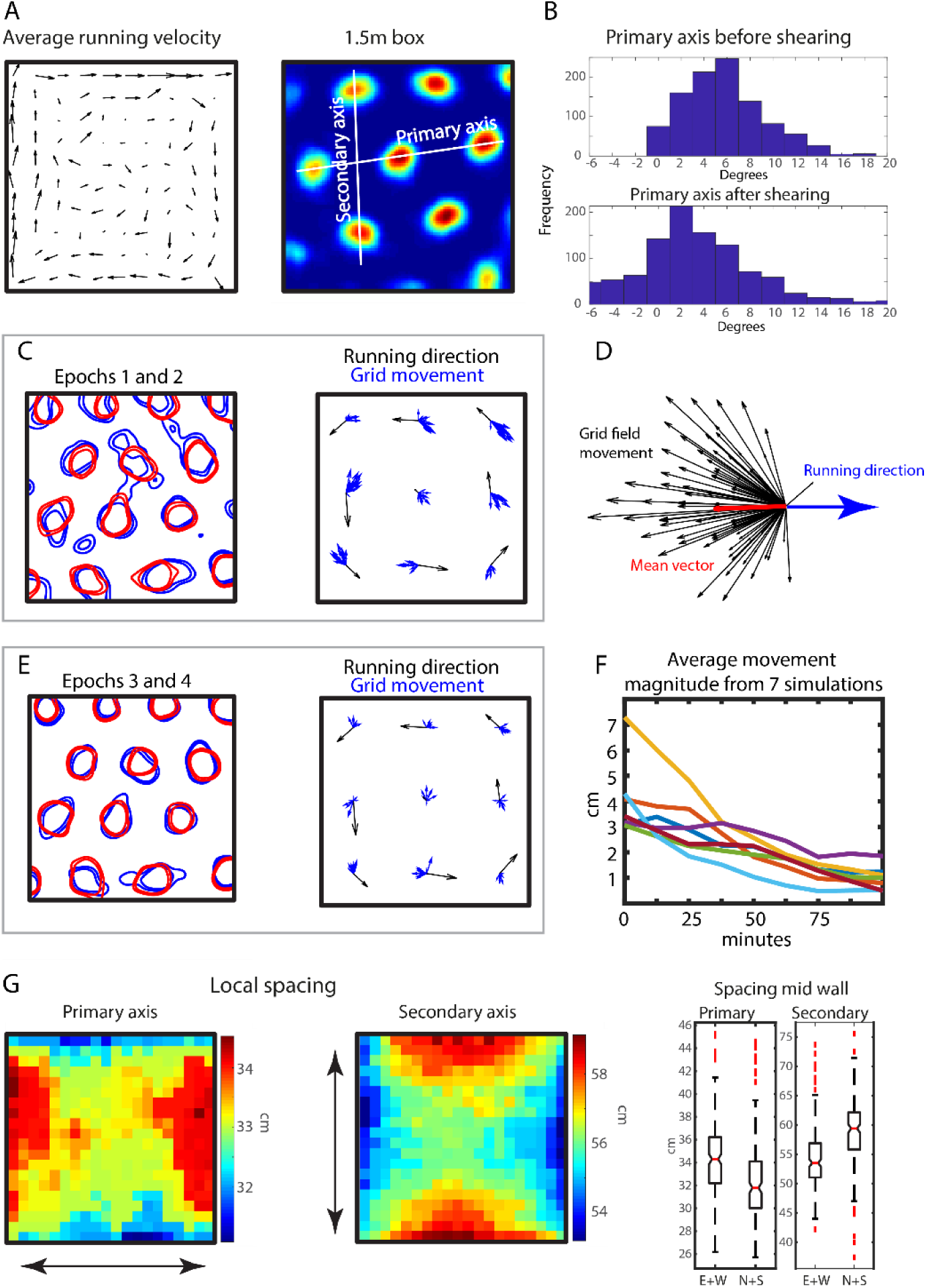
An anchoring CAN model reproduces distortions and field movement. **(A)** A CAN model with distributed anchoring was simulated with anisotropic running behavior (left) produced grid patterns (right). **(B)** The model produced a grid with an offset of the primary axis that was diminished upon minimizing the ellipticity by shearing (top vs bottom) reproducing previously described distortions. **(C)** Local cross-correlation between an initial epoch (left, blue) and a subsequent epoch (left, red), with epochs being 21 minutes long, shows that the grid fields in the model moved locally (right, 20 cells) Blue arrows show movement of individual cells and black arrows show the average running velocity. **(D)** Normalizing the movement according to the local average running direction of the simulated animal showed that the grid shift was distributed in the opposite direction from the running direction (example from C). **(E)** During later epochs the movement had subsided. **(F)** Measuring the shift every 5 minutes displays how the magnitude of the shift goes down with time (7 simulations shown). **(G)** The model reproduced local spatial distortions. The spacing of the primary and secondary axes was smaller along the walls that ran alongside those axes (arrows denote the axial direction). The boxplot displays the spacing at the mid wall of each axis (notches show the 95% confidence intervals).

Although the shift was undetectable after a few sessions, there is a possibility that movement continues albeit at a slower pace. We therefore analyzed a set of previously recorded data(Stensola et al. 2012) that spanned over weeks. Since cells were only recorded once, we here examine whether the population representation of the shape of the grid changed by measuring the offset of the grid axis that was most closely aligned to a wall of the environment. Out of 39 modules, 15 modules were recorded at more than 10 occasions. The correlation between the absolute offset of the axis and recording day showed that in 6 of these cases (40%), the orientation changed over time. In 4 out of the 6 cases, the orientation rotated towards 0° offset (r = −0.44, p = 0.025; r = −0.71, p = 0.0010; r = −0.45, p = 2.3 × 10^−5^; r = −0.30, p = 0.029) and in the remaining two cases it rotated away from 0° (r = 0.45, p = 0.01; r = 0.27, p = 0.0055).

Thus, these data show that the grid is initially volatile and moves between the first and subsequent encounter with an unfamiliar environment and that this movement occurs preferentially in the direction opposite to the animals typical running direction. This movement slows down within a few sessions so that it is undetectable by these means but on a population level the grid may continue moving over weeks.

### AN ANCHORING CAN MODEL RECAPITULATES FIELD MOVEMENT AND DISTORTIONS

The movement of the pattern suggests that the grid may change its points of anchoring. To investigate whether plastic anchoring inputs to a CAN would permit these dynamics we modelled a small CAN of 672 grid cells connected to plastic inputs from 289 place cells with place fields in a square lattice covering the entire environment. Place cells closer than 20 cm to a wall had an 20% lower firing rate than the rest to mimic the real distribution of firing rates of hippocampal CA1 place cells(Muessig et al. 2015). Each grid cell received inputs from every place cell through Hebbian synapses modelled by the BCM-rule(Bienenstock, Cooper, and Munro 1982) where the sliding threshold was modulated by the firing rate during the last 300 ms. The patterning on the cortical sheet within the CAN aligned to an edge because of the square architecture of the modeling environment. To allow for arbitrary orientations, an offset variable µ was added to the head direction. Every 100 seconds, the grid orientation was measured from an autocorrelogram of the last 300 seconds of activity and µ was updated according to the prevailing orientation. This rather artificial operation allows the CAN to slowly realign the orientation of the attractor to the pattern expressed during the last few minutes. In the brain, this operation could be performed by plastic interactions between the head direction system and the grid cells.

The balance between the collateral connections within the CAN and the inputs from the place cells was set by α so that α = 1 meant that grid cells only received inputs from other grid cells, and at α = 0, they only received place cell inputs. As α was increased, the lower inputs from the place cells made the grid smaller and less stable and at α = 1, the grid pattern was completely abolished, the firing rate was approximately 80% lower and the cells activity strongly reflected the head direction inputs (Supplementary Figure 1), similar to experimental data(Bonnevie et al. 2013). The simulated animal moved with anisotropic behavior so that whenever it reached a wall there was 0.02 higher probability per iteration of turning to the left than the right, which also leads to a higher average velocity and higher occupancy along the walls (Figure 2A). This stereotypical behavior of rats is correlated to compression distortions(Hagglund et al. 2019).

The model was run 100 times for 500 000 iterations (83 minutes) in a 1.5 m box with 289 place cells, α = 0.92, an initial position in the corner of the box and the initial orientation of the CAN at 0°. We chose a subset of 10 random cells from each simulation to produce an average that we compared between the simulations. After rotating and reflecting the grid pattern from different simulations so that their primary axis was between 0° and 15° the final pattern displayed a median primary axis offset of 5.3°, sd = 3.7 ° with few simulations with offsets close to 0° and 15° (Figure 2B, top). Minimizing the elliptic distortion by shearing the pattern along the box axis led to a smaller median primary offset of 2.9°, sd = 4.9° (Figure 2B, bottom, Wilcoxon rank test, p = 0.00010) and there was a negative correlation between the shearing factor that minimized the ellipticity and the offset of the orientation of the primary axis (r = −0.38, p = 8.3 × 10^−5^). Thus, the pattern had sheared along the N-S axes, similar to real grid cells (Stensola et al. 2015).

The development of the pattern was measured by performing a local cross-correlation in each of 9 sub-compartments between an initial epoch (Figure 2C, blue) and a subsequent epoch (red), with epochs being 125000 iterations (21 minutes) long. The grid had moved in directions that were highly distributed overall (Figure 2C right shows an example simulation, red arrows denote the shift of individual cells, 20 cells plotted), but when adjusting the shifts for the average running velocity in each sub-compartment, the shift of the grid was directed in the opposing direction (Figure 2D shows the example in C, for the whole population the circular mean direction was 167.5°, MVL = 0.63, V-test_180_, p < 10 × 10^−20^, mean shift magnitude = 3.3 cm). The deviation from 180° was smaller if half of the simulations were reflected so that the running anisotropy direction was unbiased (circular mean direction 179.5°, MVL 0.35, V-test_180_, p < 10 × 10^−20^). Although larger than in the data, the modest MVL suggest that the grid pattern also here moved in other directions. The shifts were most conspicuous in the first epochs and later subsided (Figure 2E, shows epoch 3 and 4, and Figure 2F shows the average magnitude of the shifts for 7 different simulations with measurements starting every 5 min of the duration of the simulation) but had not disappeared at epoch 3 vs 4.

To further elucidate the effects of anchoring on the finer spatial details of the grid we ran the model in a larger 2.2 m environment with a smaller spacing and a lower α to uncover whether anchoring may also reproduce local spatial compression distortions found in such an environment(Hagglund et al. 2019). 100 simulations were run for 750 000 iterations (125 min) with 625 place cells and α = 0.85. To relate the grid to both environmental axes we defined a secondary axis of the grid as the line that passes through the 2^nd^ and 6^th^ field of an autocorrelogram when counting counter clock wise from 0° and runs orthogonal to the primary axis (Figure 2A, right). Local grid spacing was measured from the 4 out of 9 equal square sub-compartments that abutted each mid-wall section. The average autocorrelogram of 10 randomly chosen cells from each simulation showed that the primary axis was compressed by 8% along the walls that aligned to the primary axis (Figure 2G; spacing difference between the E-W and the N-S mid wall bin 2.3 cm, Wilcoxon’s test, p = 8.2 × 10^−12^). The spacing along the secondary axis was similarly compressed by 12% along the walls that aligned to the secondary axis (Figure 2G bottom left and right; difference between N-S and E-W mid wall bins 5.8 cm, Wilcoxon’s test, p = 2.3 × 10^−19^). Furthermore, the grid displayed diagonal symmetric local distortions which have been shown to follow from compression distortions (Supplementary Figure 2A).

Disabling the differential place cell firing rates did not remove the compression distortion (primary and secondary axis compressed by 9% and 7% respectively, Wilcoxon’s test p = 1.2 × 10^−20^ and p = 7.3 × 10^−17^ respectively) but it did abolish the diagonally symmetric form of the grid (Supplementary Figure 2B). However, as the BCM-rule may introduce intrinsic grid formation on the single cell level independently from the CAN(Stepanyuk 2015) we speculated that the compression distortion may stem from border fields not being constrained by fields lying outside the box (Supplementary Figure 2C). The compression stemming from this effect should thus be reduced by lowering the impact of the plasticity by increasing α (and lowering the speed gain to compensate for the smaller spacing stemming from the smaller anchoring input), which is exactly what we saw. With α = 0.96 (as high as possible without losing stability of the grid) and v_gain_ = 0.018, primary and secondary axes were less compressed (both by 4%; smaller than simulations run at α = 0.85, Wilcoxon’s test p = 0.0030 and p = 4.5 × 10^−5^ for the primary and secondary axes respectively). Shearing depended on both the anisotropic behavior and the compression distortion and was abolished if either one was not present (Supplementary Figure 2D; without anisotropic behavior, median primary axis offset increased from 3.5°, sd = 3.4° to 5.5° sd = 5.1° after shearing, Wilcoxon’s test, p = 0.013; correlation between shearing factor and offset, r = −0.21 and p = 0.02; with homogenous place cell firing rates and α = 0.97 (as high as possible without losing stability of the grid), v_gain_ = 0.018, median primary axis offset went from 1.8°, sd = 3.7° to 4.9°, sd = 4.4° after shearing, Wilcoxon’s test, p = 0.0014; correlation between shearing factor and offset, r = −0.10, p = 0.28).

Thus, the model reproduced the field movement seen during novelty by re-anchoring individual grid fields to new place cells in the opposite direction of consistently traversed paths. Moreover, both shearing and compression distortions were found that depended on anisotropic behavior, heterogenous place cell activity and the influence of plasticity on single cell level pattern formation.

### THE MODEL PRODUCED TOPOLOGICAL DEFECTS IN THE PATTERN

A subset of simulations contained notable local inconsistencies in the pattern. The coherence of the grid was therefore investigated by estimating the local hexagonality of the pattern. The de-elliptified gridness score(Yoon et al. 2013) was measured locally using a sliding window autocorrelogram approach. The de-elliptified gridness score is a metric that fits an ellipse on the 6 innermost peaks of the autocorrelogram (excluding the center peak) and then transforms the autocorrelogram so that the peaks lie on a circle. The gridness score(Langston et al. 2010) is subsequently measured throughout the environment to provide an estimate for hexagonality that is not biased by ellipticity. Henceforth we will only use this de-elliptified gridness score.

Many maps exhibited a homogenously distributed high gridness score throughout the environment (Supplementary Figure 3, top) In some simulations there was a local region where the score was either below zero or undefined because the 6 innermost peaks of the autocorrelogram did not form an ellipse (Figure 3A). In the same region as these low gridness scores, the local spacing and orientation of the grid underwent a spatial phase transition around a nexus, where one or two of the axes of the grid suddenly jumped to a different configuration (Figure 3B and 3C). This jump signified that the two peaks (autocorrelograms are 180° rotationally symmetric so peaks come in pairs) closest to the center of the autocorrelogram became farther away than another pair of peaks. Thus, we concluded that in the location of the anomaly, one row of fields locally exhibited a larger spacing than a neighboring row, but farther from the anomaly the pattern was again coherent. Taken together, these transitions are consistent with a phenomenon known as a dislocation.

**Figure 3.**
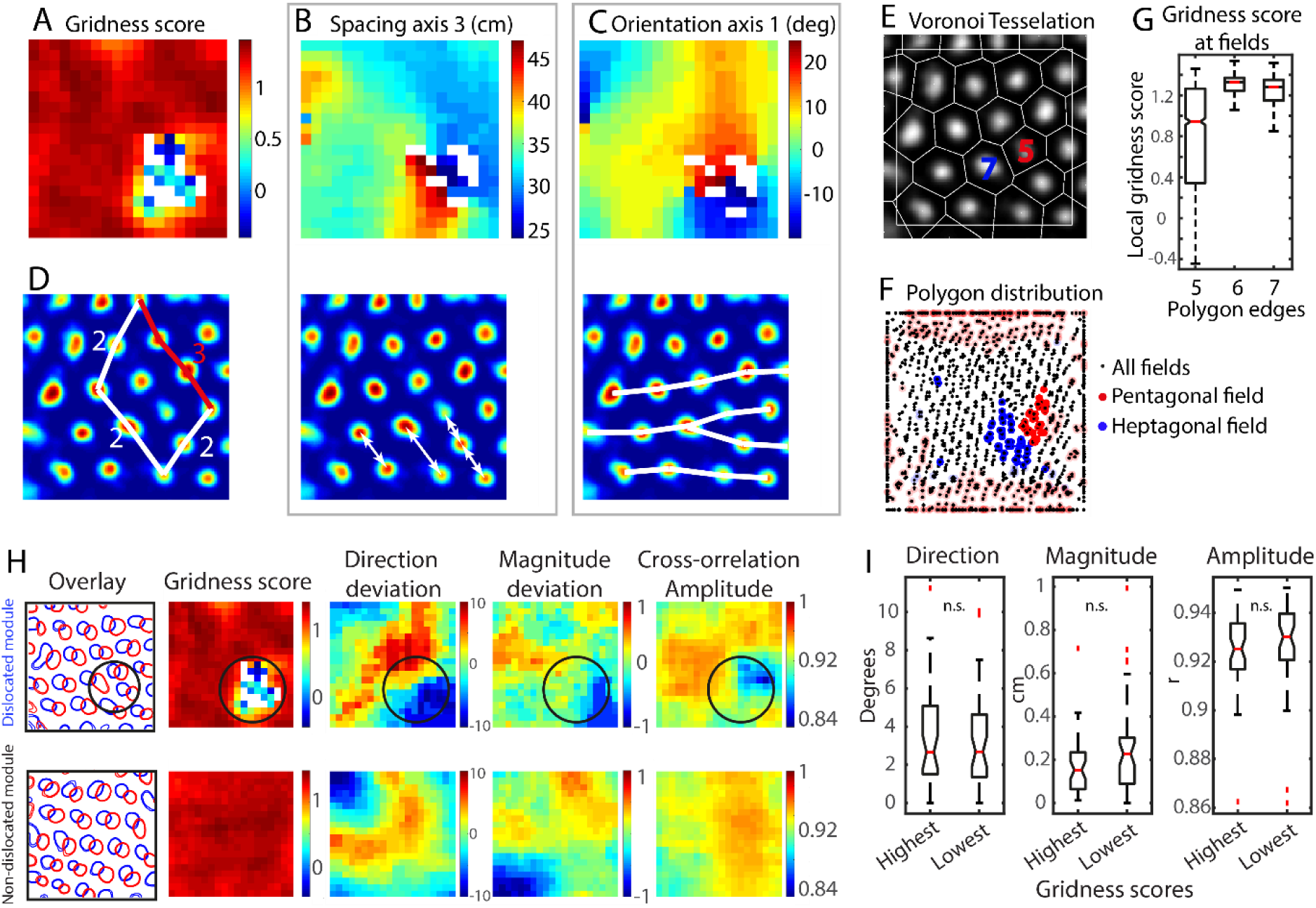
The model produces grids with topological defects. **(A)** Some simulations contained local dips in local gridness score (blue) and areas where the gridness score could not be defined (white pixels). **(B and C)** The regions of low gridness score were accompanied by a local phase transition in spacing and orientation respectively showing that these grids had a dislocation defect. **(D)** A Burgers circuit can be drawn that encapsulates the defect and shows the discrepancy in positional order. **(E)** Voronoi tesselation analyses was used to identify pentagonal and heptagonal fields. **(F)** There was a local pentagon/heptagon hot-spot in the same area as the low gridness score (same simulation as in A, B, C and D). Transparent red and blue fields denote those fields that had one Voronoi edge closer than 20 cm to a wall. **(G)** The gridness scores at the areas with pentagons, hexagons and heptagons shows that non-hexagonal Voronoi cells display a lower gridness score (outliers not shown). **(H)** The translational symmetry was examined by locally cross-correlating rate maps from pairs of grid cells (left). A dislocated simulated grid (top, dislocated region encircled) and a non-dislocated simulation (bottom) is shown for comparison. The deviation from the average direction and magnitude, and the amplitude of local cross-correlations show that the local translational symmetry is not topologically deteriorated at the locations of the dislocation but compares to the variability seen in the non-dislocated maps. **(I)** Direction and magnitude of the cross-correlation offset deviation, and the amplitude of the cross-correlation was not significant between the area of the highest local gridness scores vs the lowest local gridness scores in simulations with a dislocation (notches show the 95% confidence intervals).

Dislocations have been studied in materials science for over 80 years as they have a large impact on the properties of metals and other crystalline materials(Burgers 1939). They are defined by the

Burgers vector, which highlights the topological irregularity by drawing a loop (the Burgers circuit, Figure 3D) around the defect and defining the lattice displacement **d** as the difference in position **R** in the dislocated lattice with the corresponding position **R**_o_ in a perfectly crystalline counterpart, after integrating over the circuit. Thus **d** = **R - R**_o_ and Burgers vector **b** is then ∮**d** = **b**. In a 2-dimensional pattern, a dislocation will always have a |**b**| = 1. Moreover, dislocations can be visualized as an extra axis inserted from one of two directions, or as a pentagon - heptagon pair adjacent to each other in the lattice. A dislocation is a break in the topology and cannot be dissolved by local restructuring neighboring bonds.

To confirm the dislocations by a different method we quantified the number of pentagons and heptagons in the pattern using Voronoi tessellation by Delaunay triangulation, with grid field locations as seeds (Figure 3E). All maps had high deviation from hexagonality along the borders (Figure 3G), which is to be expected from this method, as there are no fields outside the borders to constrain the Delaunay triangulation. Therefor we disregarded any Voronoi cells that had an edge closer than 20 cm to the border (Figure 3F, transparent colored fields). The number of bins with a zero or lower gridness score, or an undefined gridness score, correlated with the mean number of pentagons (r = 0.57, p = 6.1 × 10^−10^) and heptagons (r = 0.52, p = 2.1 × 10^−8^) in the population. Moreover, the location of the pentagonal/heptagonal fields coincided with the areas of lower gridness scores (Figure 3G; median local gridness scores where there were pentagonal fields, 0.99; hexagonal fields, 1.33; heptagonal fields 1.27; 1000 cells from 100 simulations, ANOVA, F(4, 11955) = 1114, p = 2.9 × 10^−182^, Wilcoxon’s test of local gridness scores at pentagons and heptagons vs hexagons, p = 6.8 × 10^−99^).

We next asked whether the local phase structure of the dislocated simulations displayed topological breaks in the translational symmetry between grid cell pairs, such as a reflection of the phase-to-phase coupling between cells, or a disclination defect where the rotational order is altered, in contrast to the positional order which is changed in dislocations. To visualize the variations in translational phase coupling, a sliding window cross-correlation was performed between 50 cell pairs that had an average offset distance of 25% +/-10% of the grid spacing (see examples in Figure 3H). The offset between cell pairs were normalized by the mean so that a map could be constructed displaying the local average directional and magnitude deviation as well as cross-correlation amplitude between cells in the population (Figure 3H, 3 right-most panels) showing that close to areas with low gridness score the map often underwent a local change in the phase structure, but it was not much different from the variation seen in non-dislocated simulations. In each map with a gridness score of zero or below, or where the gridness score was undefined, we measured the amplitude and deviation of magnitude and direction at the minimum and maximum level of gridness score (Figure 3I). None of these variables were different across the population (difference in deviation in direction 0.3°, magnitude 4.7 cm, amplitude r = 0.0043, Wilcoxon’s test p = 0.19, p = 0.55, p = 0.07 respectively). To further quantify the phase structure, we subdivided rate maps into 9 sub-compartments and measured the phase offset between 50 cells if their average distance were 25% +/- 10%. The results showed that in locations where the gridness score was zero or smaller or undefined the offset magnitude and the offset direction displayed mean deviations of 0.4 cm and 0.4° respectively, compared to where the gridness score was higher than 0 (Wilcoxon’s test, p = 0.35 and p = 0.018, respectively). Although significant, the phase offset deviation within dislocated regions were small and the overall translational symmetry between cells was thus mostly contained, which is not surprising since we impose the phase structure with the synaptic connectivity architecture of the CAN.

### DISLOCATIONS ARE FOUND IN REAL GRID CELLS

To investigate whether the dislocations described above could occur in real grid cells we analyzed a data set of 131 grid cells from 5 modules recorded from three animals foraging in a 2.2 m square environment(Stensola et al. 2012). Grid modules were selected from the population if they had a mean spacing of less than 60 cm and were part of a module consisting of more than 10 cells. Modules were assigned in a previous study from which the data came from(Stensola et al. 2012). De-elliptified gridness scores were measured locally throughout the environment using a sliding window with a size of 73 cm (Figure 4A). Three of the modules (here referred to as module 1-3) described a consistently high gridness score throughout the environment. However, two modules (module 4 and 5) contained a region of negative or undefined gridness score denoting low hexagonality. At these locations, local spacing and orientation was found to exhibit a similar transition as that found in the modelled grid (Figure 4C and D).

**Figure 4.**
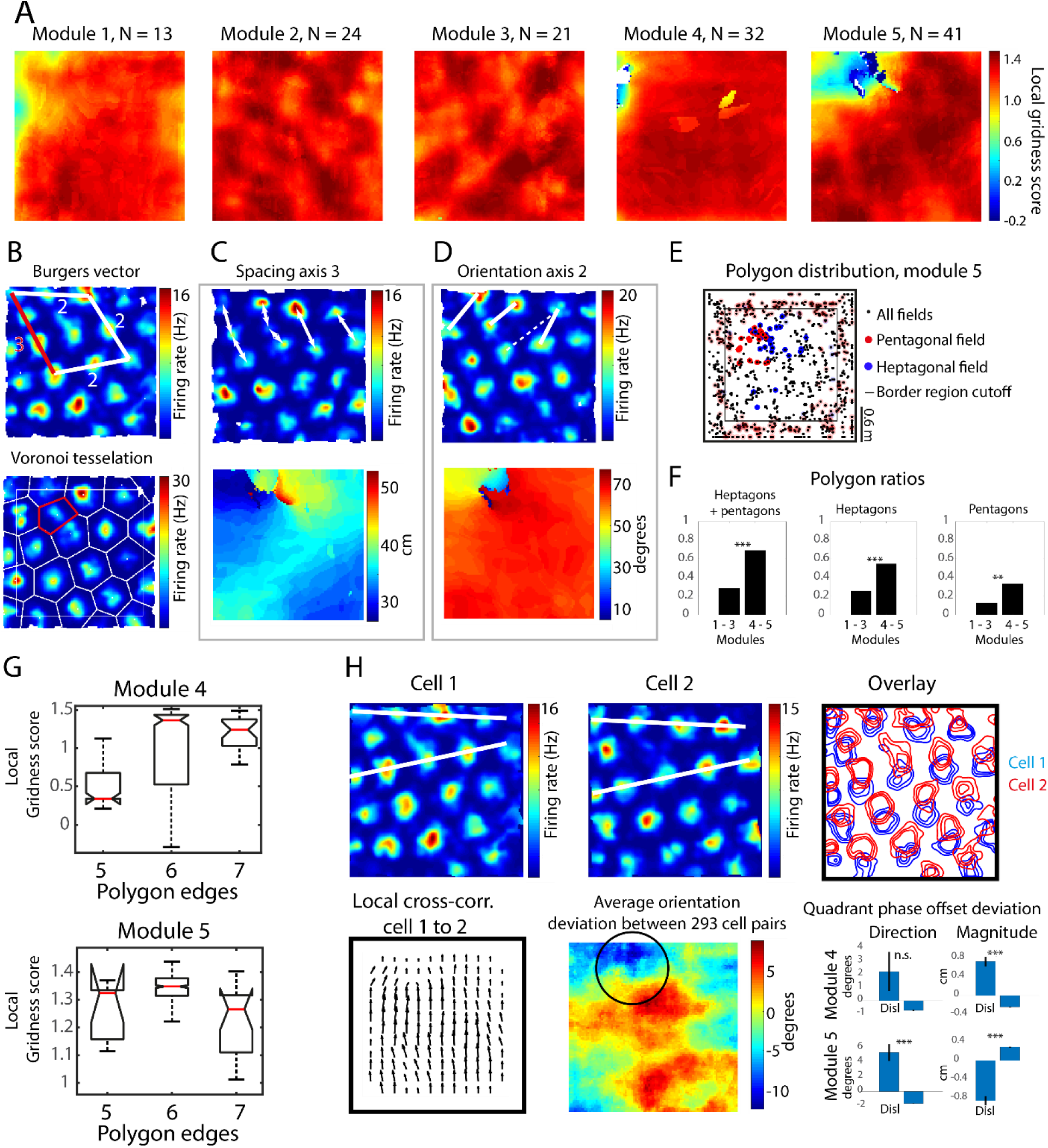
Topological defects may be found in real grid cells. **(A)** Local gridness score maps of 5 modules reveals that two of the modules contain a a region of low or undefined gridness score. **(B)** An example of how the Burgers circuit encapsulates the dislocation (top) and an example of a Voronoi diagram of a grid with a pentagonal field in red (bottom). **(C and D)** The local shape of the grid in the dislocated modules demonstrated sudden transitions visible in the local spacing and orientation maps respectively (bottom). **(E)** The distribution of polygons as measured using Voronoi tessellation (B, bottom) shows the accumulation of pentagonal and heptagonal fields in the dislocated area. **(F)** Modules 4 and 5 contained a higher ratio of pentagons and heptagons compared to the other three modules. **(G)** The local gridness scores in module 4 (top) and module 5 (bottom) were lower at pentagonal and heptagonal fields compared to hexagonal fields (outliers not shown, notches show the 95% confidence intervals). **(H)** The phase offsets between different cells in a module (H, top row two example cells) is mostly maintained also across dislocations. Local cell pair cross-correlations displays small deviations across the environment (bottom left). On the population level (mid bottom) the deviation from the mean orientation is different in the dislocated area (encircled region). The mean vector for the deviations from the mean in dislocated quadrants vs the rest (bottom right) shows that the translational symmetry is different in direction in module 4 and in both magnitude and direction in module 5. Error bars show standard error.

To investigate the hexagonality and deviations from it directly, fields were defined as regional maxima in locally normalized oversmoothed rate maps (Figure 4B, bottom) with a smoothing kernel proportional to the grid spacing (see Methods). The Voronoi method was used as described above, disregarding any Voronoi cells that had an edge closer than 20 cm to the border. Non-hexagonal Voronoi cells were found scattered throughout the environment in each module, likely false positives stemming from the noise of low sampling, individual differences in firing rate and disproportionate coverage. However, in module 4 and 5 there was a local hot-spot of non-hexagonal Voronoi cells (example in Figure 4E). For each rate map the numbers of pentagonal and hexagonal Voronoi cells were counted. Module 4 and 5 contained a higher ratio of grid cells with at least one heptagonal or pentagonal Voronoi cell (Figure 4F left; chi-square test, chi = 18.9, p = 1.4 × 10^−5^), higher ratio of heptagons (Figure 4F mid; chi-square test, chi = 11.5, p = 6.7 × 10^−4^) and a higher ratio of pentagons (Figure 4F right; chi-square test, chi = 6.4, p = 0.011). The local gridness score was lower at pentagonal and heptagonal fields compared to hexagonal fields (Figure 4G; median gridness scores at pentagonal fields for modules 4 and 5 respectively, 1.32 and 0.34; hexagons, 1.35 and 1.40; heptagons, 1.27 and 1.24, F(4, 434) = 13, p = 0.0008 and F(3, 295) = 73, p = 1.9 × 10^−35^; Wilcoxon’s test for pentagons and heptagons vs hexagons, p = 2.1 × 10^−6^ and p = 1.0 × 10^−20^ for modules 4 and 5 respectively). The local gridness score and Voronoi analyses were also applied to a data set of three modules from animals running in a geometrically manipulated environment from a previous study(Wernle et al. 2018). 2 out of 3 modules also displayed negative/absent local gridness scores and had a higher ratio of heptagons and pentagons (supplementary Figure 4).

The two dislocations in the 2.2 m data set were conveniently located within quadrant boundaries. To examine whether the grid pattern was different in the dislocated quadrants, the amplitudes of the cross-correlations within cells of the environment was measured in the four quadrants. The mean of the three innermost maxima of the cross-correlograms were used to get a consistent measure of the amplitude. The amplitude was significantly lower for modules 4 and 5 containing a dislocation (ANOVA of cross correlation amplitudes between quadrants for each cell F(5, 72) = 1.00, p = 0.43; F(5, 138) = 1.14, p = 0.34; F(5, 120) = 0.86, p = 0.51; F(5, 186) = 5.31, p = 0.00039; F(5, 240) = 14, p = 3.4 × 10^−12^ for modules 1-5, respectively). The quadrants containing the dislocation had the lowest mean cross-correlation amplitude (mean cross correlation amplitude in the dislocated quadrant in module 4, r = 0.35, sd = 0.12 vs the three other quadrants, r = 0.38, sd = 0.13, Wilcoxon’s test, p = 0. 021; and r = 0.37, sd = 0.12 in the dislocated quadrant of module 5 vs the three other quadrants, r = 0.41, sd = 0.12, Wilcoxon’s test, p = 2.0 × 10^−5^). Thus, within each rate map, the grid in the quadrant containing a dislocation had a different shape compared to the three other quadrants.

As in the model, we asked whether the local phase structure of the dislocated modules displayed any topological breaks in the translational symmetry of the grid, such as a reflection of the phase-to-phase coupling between cells or a disclination defect. Local phase deviation maps showed that similar to the model, the offset displayed a local deviation at the region of the dislocations which was not abrupt but continuous (see example in Figure 4H, top three panels and bottom left).

To quantify the local translational symmetry, each quadrant of the rate maps was cross-correlated between cells within each of the dislocated modules. The distance and direction offset from the middle to the closest peak in the cross-correlogram was measured. Local population deviation of direction and distance between cells were evaluated by subtracting the mean magnitude and direction of the offset from each measurement (Figure 4H, bottom right). In module 4, the offset magnitude between cells was larger in the dislocated quadrant (deviation from mean offset magnitude 0.73 cm, sd 3.2 vs −0.24 cm, sd 0.20 in the three remaining quadrants, Wilcoxon’s test, p = 1.0 × 10^−19^). However, the cells had the same directional offset in the dislocated region as in the rest of the environment (circular mean deviation 2.2° in the dislocated quadrant vs 0.72° in non-dislocated quadrants, Fishers test for median circular variables, p = 0.57). In module 5, the phase offset magnitude was significantly smaller (deviation from mean offset −0.87 cm, sd = 3.3 in dislocated quadrant vs 0.29 cm, sd = 0.35 in the rest, Wilcoxon’s test, p = 5.6 × 10^−31^) and the directional component was different in the dislocated quadrant (circular mean deviation in orientation 5.4°, circular sd 38.2° in the dislocated quadrant vs −1.7°, circular sd 34.2, Fishers test for median circular variables, p << 0.001). Although significant, these results show that the variability in phase relationships between cells is small in a dislocated region. The translational symmetry between cells is thus not topologically disturbed but may stay roughly consistent also across dislocations.

## Discussion

We have shown that the layout of the grid pattern is experience dependent and may be in a state of change depending on the behavior during initial encounters of an environment as well as over longer time spans. The novelty induced re-anchoring could be reproduced by a CAN that learns to anchor to place cell inputs. Moreover, the learning CAN explained previously described spatial distortions and predicted topological defects with retained translational symmetry between cells. These topological defects were found in a data set of animals running in a large environment.

Grid cells have been shown to fire in anticipation of grid fields(Kropff et al. 2015; De Almeida et al. 2012). This effect may lead to associative re-anchoring during times of high plasticity that likely occurs during an encounter with a novel environment. In our model, upon entering a grid field, a grid cells firing rate increases which leads to potentiation of place cell synapses from place cells with place fields at this grid field entry. However, potentiation according to adapting Hebbian learning, such as with the BCM-rule used here, saturates with time, which means that when the animal exits the grid field place cell synapses from place cells with fields at the exit point will be less potentiated or may even be depressed. Thus, grid cells slowly shifted their representation by changing which place cells they anchored to. Within session movement was not analyzed because of a lack of data. Although possible, we find it less likely that the field shift occurred solely during the inter-session rest period and more likely to be a gradual process unfolding during a session. The possible influence of top-down inputs from hippocampal cells do not change the likely scenario that grid cells also influence and stabilize place cell fields in an interactive process (Renno-Costa and Tort 2017).

Other evidence of local shifting and re-anchoring overriding the coherence of the grid pattern have been found where grid fields moved locally close to reward sites(Boccara et al. 2019), home locations(Sanguinetti-Scheck 2019) or where the environment had been geometrically altered(Krupic et al. 2018; Wernle et al. 2018) further implicating the behavioral impact of local anchoring on the grid pattern. Here we found a close connection of the grid movement to the action of the animal suggesting that the previously described grid field changes may be accounted for by the behavioral constraint that the rearranged environments produced in contrast to stemming from direct sensory inputs.

Several anchoring CAN models have been suggested where the grid has been proposed to be reset by borders(Hardcastle, Ganguli, and Giocomo 2015; Ocko et al. 2018; Keinath, Epstein, and Balasubramanian 2018), salient visual landmarks(Mulas, Waniek, and Conradt 2016), landmark cells(Campbell et al. 2018). In the current work however, anchoring stemmed from a distributed population of place field inputs, akin to(Guanella, Kiper, and Verschure 2007; Kropff and Treves 2008; Dordek et al. 2016; Stepanyuk 2015). Both the novelty induced grid field shifting and the dislocations depended on place cells with place fields far removed from any immediate salient cues. The most likely candidate for such an input stems from the CA1, which sends a massive descending input to the deep layers of the MEC. Other possible sources include the subiculum and a population of place cells that reside in the retrosplenial cortex, since both these areas also project heavily to the MEC. Place cells may form fields attached to landmarks and salient local features but are to a great extent found in the open field of an experimental box(Muessig et al. 2015).

Local compression distortions and shearing could be explained by anisotropic behavior and the intrinsic periodicity induced by the synaptic plasticity(Stepanyuk 2015). The latter may depend on a lack of traversal of fields lying outside the environment which leads to a lesser force separating the fields. The lack of constraining neighboring fields along the secondary axis would impose a stronger force than along the primary axis which is why the secondary axis is relatively more compressed than the primary axis. This effect may further explain why the pattern is not only rotated by the anisotropic behavior but shears because of unequal compression between the primary and the secondary axis.

Dislocations in the grid system have been suggested as a proof of continuous attractor dynamics, if the translational symmetry between cells are preserved(Burak and Fiete 2009), which is what we found. However, a CAN with a dislocation in its cortical sheet could not be stable over time which is a strong evidence for an external input. Furthermore, the fact that dislocations could occur in the open field, removed from proximal cues and without environmental manipulations, is a strong evidence for distributed anchoring not dependent on direct sensory input from landmark or border cells. Dislocations form an entanglement in the spatial attractor landscape formed within the hippocampal formation which may have cognitive and behavioral consequences. Traversing the dislocation suddenly changes the spacing and orientation of the grid possibly providing a source of confusion or less fidelity of the representation of space. Dislocations formed serendipitously in the model and, although not addressed directly in the current study, may be biased by several factors. Large environments, small grid spacing, and strong anchoring could all lead to a decrease in long range interactions of the pattern and subsequent breaks in the topology of the grid. Possibly they are more prone to form from two or more disparate anchoring points between which the pattern could not coherently consolidate, which may predispose them in geometrically altered environments (Supplementary Figure 4). If and how dislocations may generalize to other attractor systems of the brain is an intriguing question for future studies.

Taken together, these experimental and numerical results suggest that the grid may be anchored by distributed place cell inputs. Such anchoring can explain why grid fields move in opposition to animals typical running pattern during learning. Moreover, local anchoring to place cells and stereotypical behaviors may explain the origin of elastic distortions and topological defects in the grid pattern.

## Methods

### ELECTROPHYSIOLOGY AND SURGERY

The experiments were performed in accordance with the Norwegian Animal Welfare Act and the European Convention for the Protection of Vertebrate Animals used for Experimental and Other Scientific Purposes. Five male Long Evans rats were implanted with a multi-tetrode device carrying 4 or 14 tetrodes atop the medial entorhinal cortex (MEC) at 4.7 mm left of bregma and 0.5 mm rostral to the transverse sinus. Recordings were made using the Neuralynx (Neuralynx, Tuscon, AZ) or Axona (Axona Ltd., UK) acquisition systems. Spikes were sorted manually using the MClust software (A.D. Redish). Tetrodes were lowered over days or weeks until strongly theta modulated neurons were found signifying the location of the MEC. Animals were trained to forage for cereal crumbs in a 1.5 m box. LEDs on the device was tracked by a recording camera situated in the ceiling. When multiple grid cells were stably recorded in a familiar box the animal was subjected to a novel 1.5 m square, 2.4 m triangular or 1.8 m triangular enclosure in a novel room or novel box with a different geometry in a familiar room where cells were continuously recorded for several sessions.

### DATA PROCESSING AND ANALYSES

All data analyses and modelling were performed using MATLAB (Mathworks Inc, MA). Grid cells were determined by repeated shuffling of the experimental data as previously described (Langston 2010). If the gridness score was larger than that of the 95th percentile of the grid scores from the shuffled data, the cell was defined as a grid cell.

Rate maps of well separated neural signals were produced as described earlier(Sargolini et al. 2006). Position estimates were smoothed with a 15-point median filter. The position data were sorted into 1.5 cm bins and a smoothing algorithm using a Gaussian kernel with a sigma of 6 cm was applied to the spike position data, which was normalized by the spatial occupancy. Locations further than 3 cm away from the animal’s path were considered unvisited. The local shift of the grid from one session to the next was measured by subdividing each map into 9 equal bins and measuring the offset by cross-correlation. Average running velocity was also collected from each sub-compartment and the grid shift was normalized to the average running direction by subtraction.

Parameters of the grid were measured in autocorrelograms where the location of the six innermost local maxima, excluding the middle peak, were extracted. The mean orientation of three axes of the grid in each module was measured and a primary axis was determined as the axis that was closest to either the N-S or E-W axis of the recording enclosure. All cells within a module/simulation were rotated and/or reflected consistently depending on the mean orientation of the primary axis of each module/simulation so that it ended up between 0° and 15° offset from the E-W axis.

To evaluate the ellipticity of the grid, an ellipse was fitted to the six innermost peaks of autocorrelograms (excluding the middle peak) using a least-square criterion and the orientation of the long axis of the ellipse was defined within the interval 0° to 180° (Fitzgibbon 1999). Ellipticity was defined as the length of the semi-major axis divided by the semi-minor axis.

In the shearing analyses, the position of the six innermost peaks was incrementally sheared by step-wise increasing the shearing factor *q* in

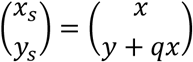

along the y axis and the value of the shearing factor *q* and primary axis offset was measured where shearing produced the lowest the ellipticity.

To measure the de-elliptified gridness score, autocorrelograms were rotated to align their elliptic orientation with the environmental orientation after which they were compressed along this axis to retain a circular layout of the innermost 6 peaks. After this transformation the gridness score was measured (Langston 2010).

Local grid parameter maps were constructed by sliding a 74 × 74 cm window across 10 randomly chosen cells from each simulation, calculating an autocorrelogram at every 7.5 cm. Autocorrelograms were averaged between cells to get a mean representation of the grid at each location of the environment. Local spacing (Figure 2G, 3B, 4C), orientation (Figure 2B, 4D, Supplementary Figure 2) and gridness scores (Figure 3 and 4) was measured in each simulation and was then averaged to produce the final maps. However, in the data analysis all cells from respective modules were used and the step-size of the sliding window was 1.5 cm.

Local phase deviation maps were constructed by a sliding window cross-correlation in steps of 4.5 cm or 1.5 cm per measurement in the model and data respectively. In the model, 50 randomly chosen cell pair combinations from each simulation was cross-correlated locally in a 74 × 74 bin window if their grid offset was between 15% - 35%. At each location the peak closest to the middle was interpreted as the translational phase offset between the cells. The mean offset was measured from the full map cross-correlation which was subsequently subtracted from the local maps to produce maps describing the local deviation from the mean. Magnitude, direction and amplitude of the cross-correlation offset peak was then extracted and plotted as individual parameters in color coded maps. In the data the process was the same except for the number of cell pairs used (293), the size of the window (30 × 30) and the step size = 1.5 cm.

### GRID POLYGONS

Fields were defined as local maxima in rate maps produced by two filtering steps. Rate maps were first locally normalized by being divided by a gaussian over-smoothed version of themselves with a *σ* = 15 cm. Next, in the model data the resulting map was further smoothed with a *σ* = 9 cm. In the real data the *σ* for the second smoothing was

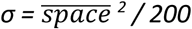

derived from the average spacing 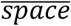of the entire module. Field locations were extracted as local maxima and used as seeds for Delaunay triangulation. Grid field polygonal structure was analyzed by Voronoi diagrams derived from the Delaunay triangulation.

### MODEL

The model consists of a periodic CAN with 24 × 28 grid cells situated on a twisted torus(Guanella, Kiper, and Verschure 2007), connected to 289 or 625 gaussian place cells in the 1.5 m and 2.2 m boxes respectively, distributed evenly every 9 cm with a sigma of 15cm. Each grid cell integrates grid cell inputs, place cell inputs and velocity input, and after this the synaptic weights from the place cells are updated according to the BCM rule with a temporal sliding threshold β based on the average firing rate for each neuron for the last 300 ms. The balance between the place cell and grid cell inputs to a grid cell is set by α.

The firing rate *g* of neuron *i* at time *t* is

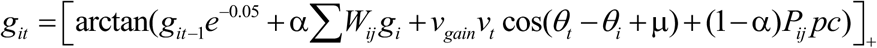

Where

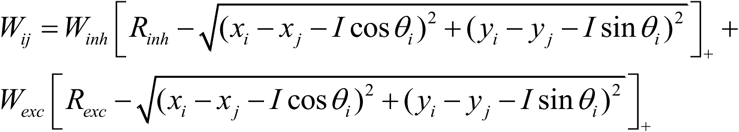

are the weights within the CAN. […]_+_ is the Heaviside threshold-linear function, *R*_*inh*_ = 10 and denotes the radial extent of the inhibitory connectivity = and *R*_*exc*_ = 3 is the radial extent of the excitatory connectivity, *W*_*inh*_ and *W*_*exc*_ is the strength of the inhibitory and the excitatory connectivity set at −0.02 and 0.05 respectively. Thus, grid cells are connected by excitatory synapses to close neighbors and inhibitory synapses to distant neighbors as inferred from functional connectivity and noise correlation studies (Tocker, Barak, and Derdikman 2015; Dunn, Morreaunet, and Roudi 2015). *x* and *y* denote positions on the cortical sheet, and *I* is an offset of 2 neurons, shifting the activity of this neuron towards the preferred direction of θ_*i*_ (one of the 4 directions (N, W, S or W), neurons with different preferences are tiled evenly over the cortical sheet). *v*_*gain*_ = 0.0023 and 0.0032 is the speed gain in the 1.5 m and 2.2 m environments respectively unless otherwise noted, *v*_*t*_ is the current speed, θ_*t*_ is the current running. *µ* is measured as the average orientation of the three grid axes within (−90,90] from an autocorrelogram of the grid produced during the last 300 s of activity and is measured from 10 grid cells every 100 s. The whole expression saturates with the inverse tangent function and cannot be negative.

*P*_*ij*_ denotes the weights from place cell *j* to grid cell *i* with *pc*_*j*_ being the firing rate of place cell *j. α* sets the gain and balance between place cell and grid cell inputs with *α* = 0.92 in the 1.5 m box and 0.85 in the 2.2 m box unless otherwise noted. Weights were initialized as a normal distribution centered on 0.1 with a *σ =* 0.003 which after normalization leads to a distribution with a peak around 0.6.

*P* is modified using the BCM-rule as

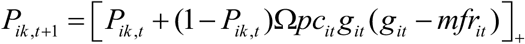

and normalized similar to (Kropff et al) with the Euclidian norm so that

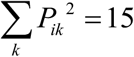

Thus, the synaptic weights have a soft upper boundary approaching 1. *mfr*_*it*_ is the temporal average activity of neuron *i* over the last β = 300 ms, and *Ω* = 0.2 is the learning rate.

### RUNNING BEHAVIOR

The simulated animal moved along a random path with speed *v* according to

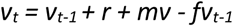

and position updated as

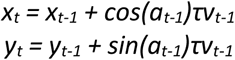

with

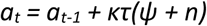

where *f =* 0.01, *τ* = 0.015s, the turning rate is *κ* = 9.2, *r* and *n* are drawn randomly from a standard distribution, *ψ* is 0 unless encountering a wall when it denotes the probability of turning left = −0.02, *mv* = 30cm/s sets the median speed. The speed was threshold at 5 cm/s.

Model grid cell ratemaps were produced as a 99×99 (1.5 m box) or 147×147 (2.2 m box) binned 2d histogram of firing rate divided by the occupancy.

## Acknowledgements

We thank Edvard I Moser and May-Britt Moser for insightful comments on the manuscript as well as for giving their consent to use these data, and Benjamin Dunn for discussions on the model. Moreover, we thank A. M. Amundgård, K. Haugen, E. Kråkvik, and H. Waade for technical assistance.

---

**Supplementary figure 1.**
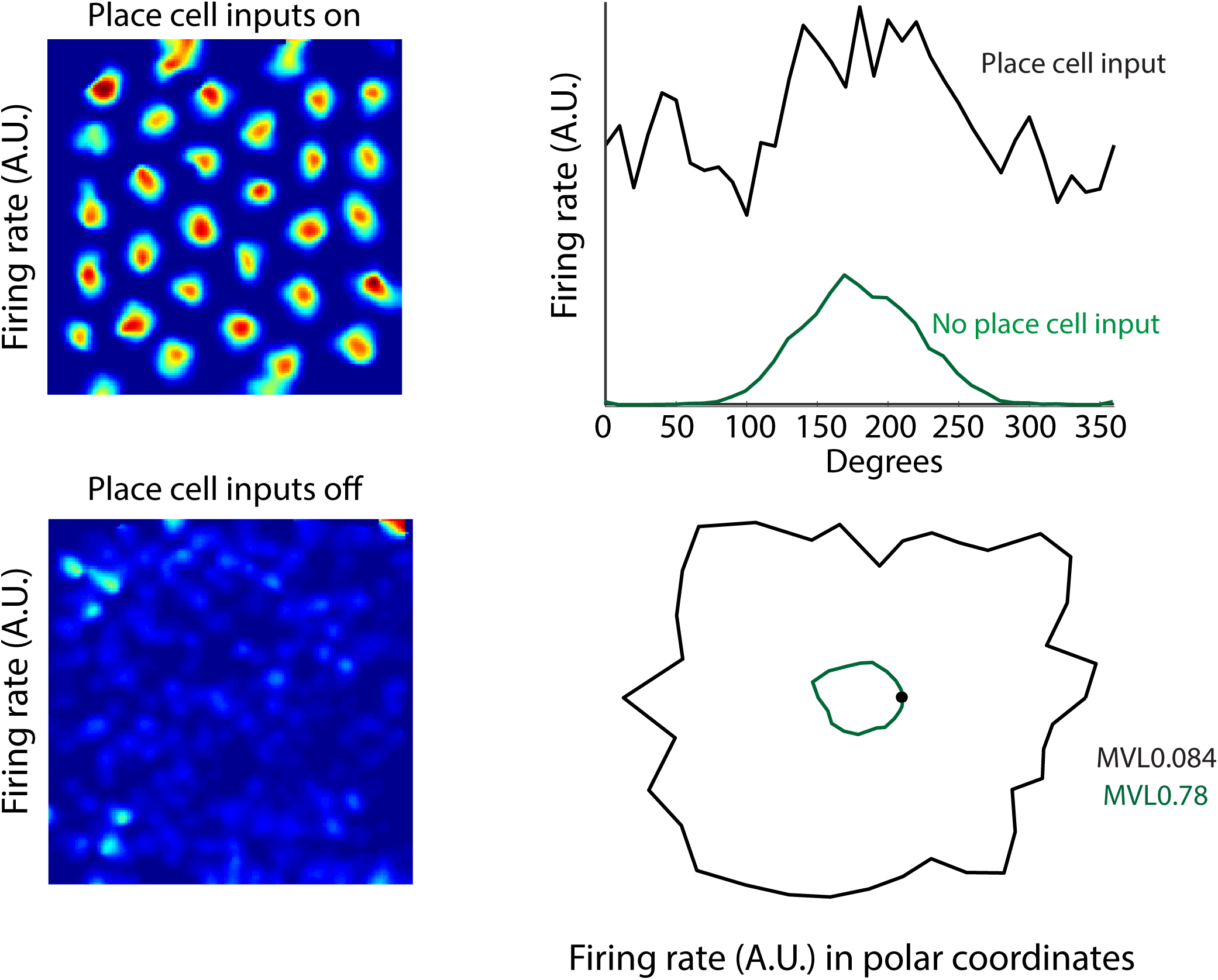
Removing the place cell input to the model abolished grid cell activity (lower left) and revealed a strong head directional component (right, green).

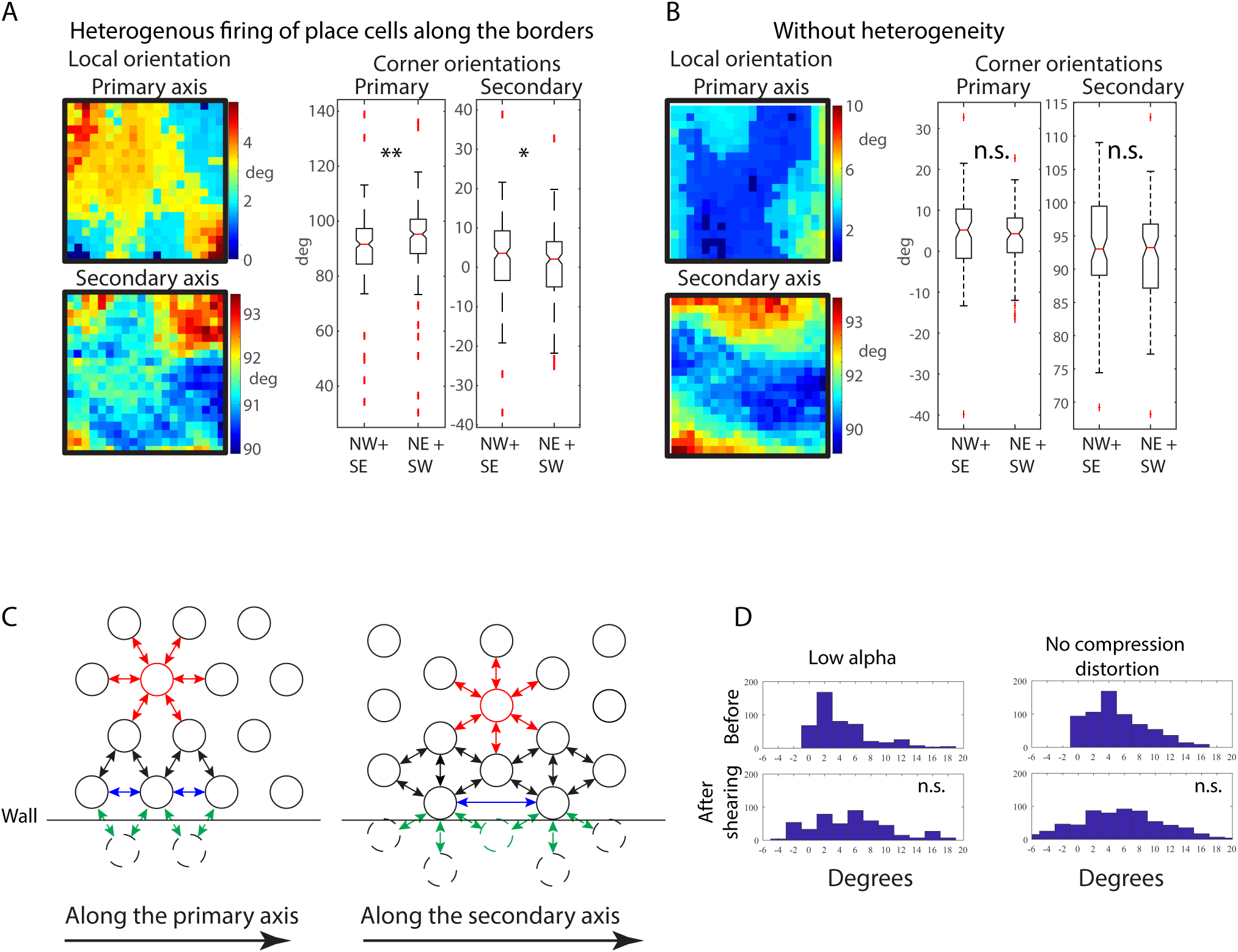
A. Diagonally opposing corners displayed similar orientations, both of the primary and the secondary axes but different from the other pair of corners (median orientation of the primary axis in NW + SE corner 1.6° and NE + SW 3.8°, Wilcoxon’s test, p = 0.0045. Median orientation of the secondary axis in NW + SE corner 93.0° and NE + SW 91.9°, Wilcoxon’s test, p = 0.026). This diagonal symmetry is a hallmark of the border compression distortion. The notches in the boxlot show the 95% confidence intervals. B. Disabling the differential place cell firing rates eliminated the diagonal symmetry (median orientation of the primary axis in NW + SE corner 5.2° and NE + SW 3.8°, Wilcoxon’s test, p = 0.20. Median orientation of the secondary axis in NW + SE corner 93.0° and NE + SW 92.2°, Wilcoxon’s test, p = 0.19). The notches in the boxlot show the 95% confidence intervals. C. Fields in the middle of the environment (red) are constrained from all directions. Fields along the borders lack neighboring fields outside the box (dashed circles) and are thus less constrained (green arrows denote lacking forces) meaning the axis along the wall may compress (blue arrows). The lack of constraint has a stronger effect along the secondary axis since the lacking border field (green dashed circle) would have had a larger constraining influence on the spacing. D. The grid did not shear if there was no anisotropic behavior (left) or if there was no compression distortions because of a homogenous place cell firing rates and a low α (right).

**Supplementary figure 3.**
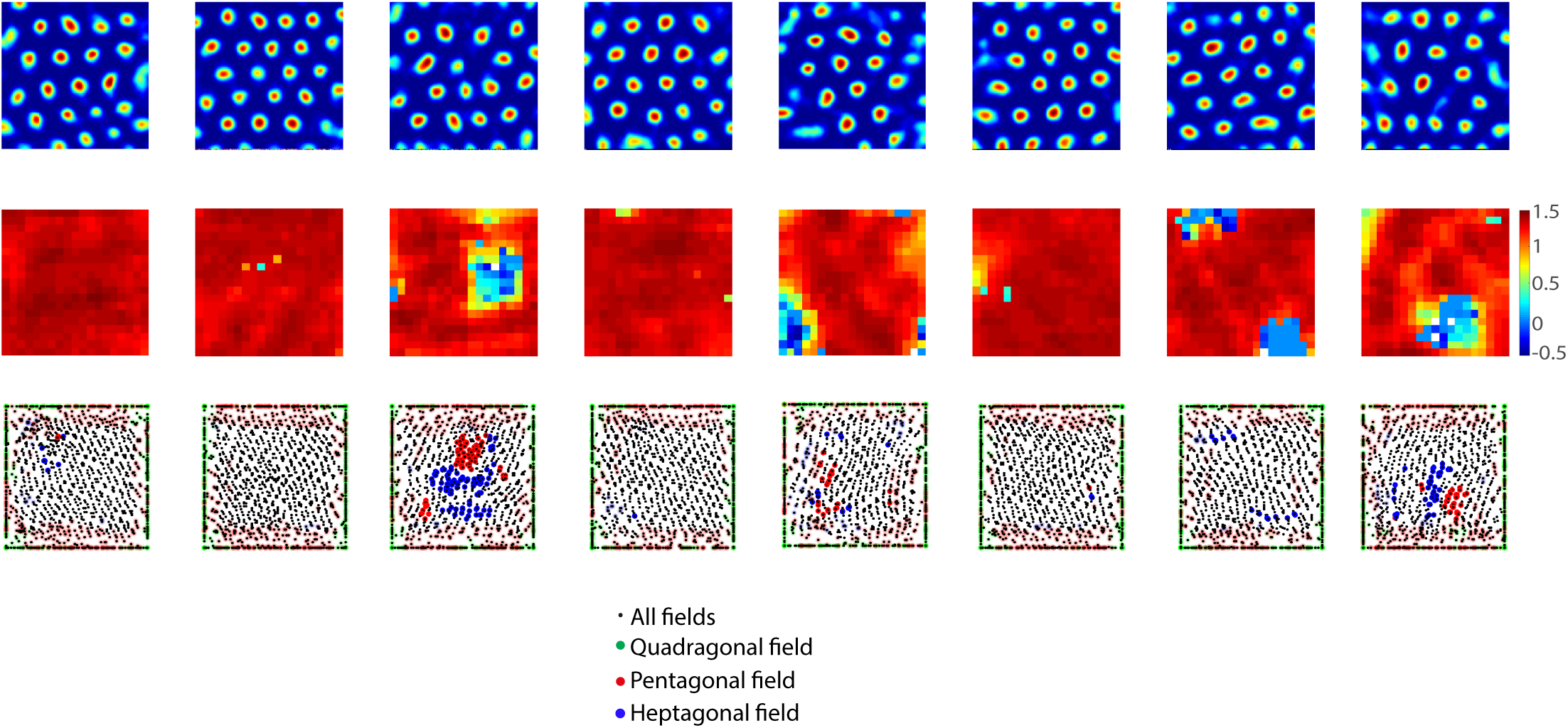
Example rate maps, local gridness scores and Voronoi diagram summaries from the 2.2 m simulations. The Voronoi maps shows the fields of 50 cells in each map (black dots) and deviations from hexagonality as red (pentagons), blue (heptagons) or green (quadragons). Transparent fields have an edge of their Voronoi cell lying outside of the cutoff of 20 cm from the wall.

**Supplementary figure 4.**
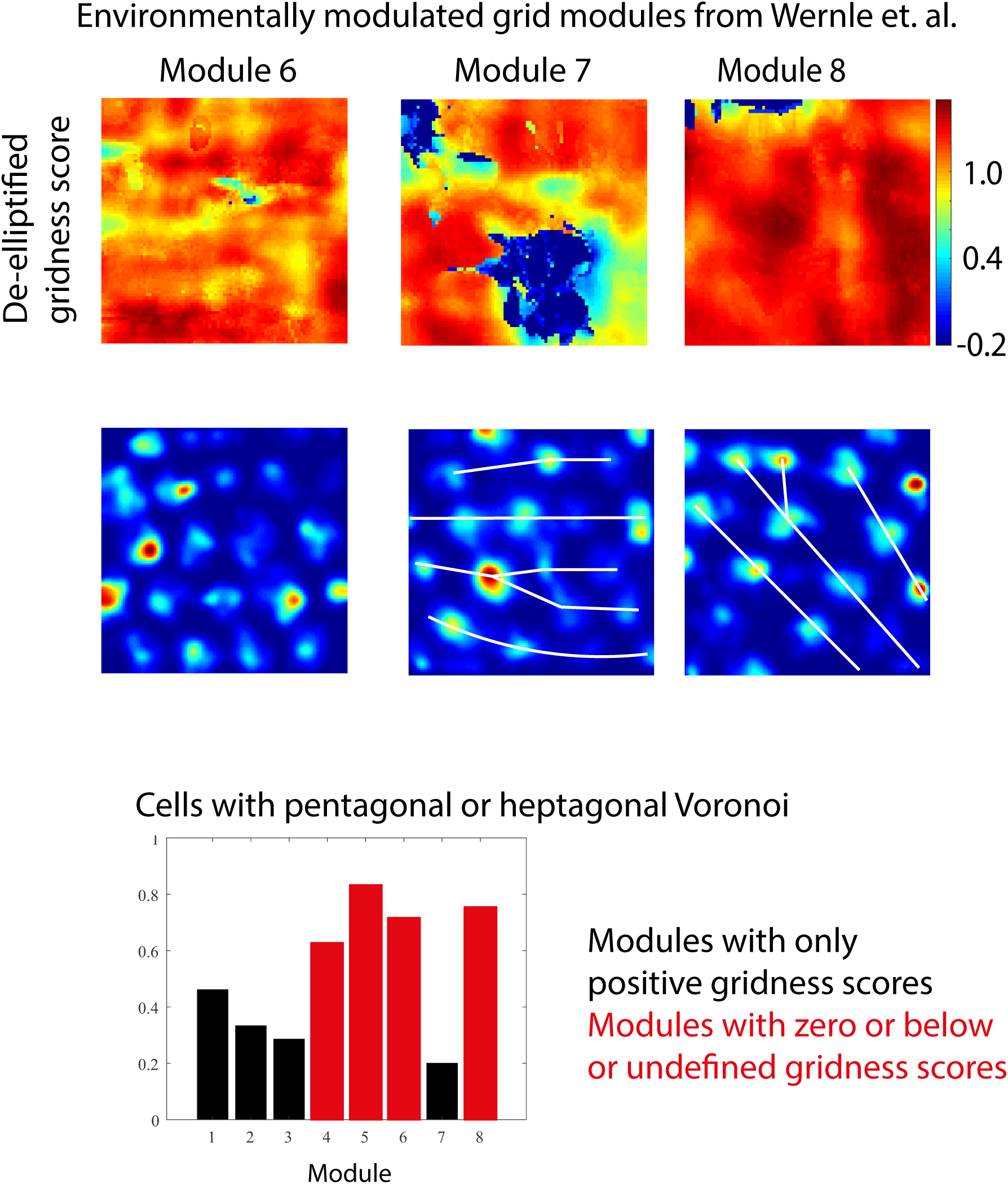
A dataset where animals had been subjected to a merging of two 1×2 m familiar environments was also investigated (Wernle et al. 2018). From 13 animals, three grid modules had more than 3 cells with a grid spacing less than 60 cm. Local gridness score maps (top) revealed that two of the modules contained areas of low fidelity, and in one of the (Module 7) seemed to contain 2 such areas. The ratio of pentagonal or heptagonal Voronoi cells is significantly higher in the modules that had a zero or below or undefined gridness score (chi = 22.4, p = 2.2e-6).

